# Identification and performance of environmentally-driven recruitment relationships in state space assessment models

**DOI:** 10.1101/2025.07.01.662400

**Authors:** Gregory L. Britten, Elizabeth N. Brooks, Timothy J. Miller

## Abstract

Environmentally-driven recruitment relationships are important for understanding fisheries’ responses to climate change; however they are difficult to estimate due in part to large variability in the recruitment process. State space models provide a promising way forward in allowing the characterization of multiple sources of stochastic variability. Here we conducted a large simulation-estimation study using environmentally-driven recruitment relationships to evaluate the effects of operating model characteristics on state space assessment model performance. We generally find low parameter and assessment bias across operating and estimating model combinations; however, some assessment and parameter bias is present under conditions of high recruitment variability and low spawning stock biomass contrast. Model identifiability for the correct functional form of the stock recruit relationship and the environmental relationship was generally poor, and projections were insensitive to assumed values of the environmental driver. We recommend the use of random effects on recruitment in state space assessments and caution against explicit stock-recruitment relationships. We encourage future work on environmental nonstationarity, which will be of increasing importance as exploited fish stocks experience accelerating rates of environmental change.

## Introduction

Recruitment refers to the number of fish entering the harvestable population per year and has been a focus of fisheries research since Hjort’s pioneering work over 100 years ago [*Hjort*, 1914]. Despite significant research effort, accurately predicting annual recruitment (i.e. the ‘recruitment problem’) has proved difficult, leading fisheries scientists to question whether recruitment should be modeled as a function of the spawner population or whether stochastic factors dominate [*Barrowman and Myers*, 1996; *Myers*, 1998]. The recruitment problem presents a continual challenge for fisheries stock assessment which is tasked to set sustainable harvest limits and evaluate whether a stock is overfished or experiencing overfishing.

Several environmental processes have been linked to recruitment variability, including physical transport of eggs and larvae [*Bolle et al.*, 2009; *Boucher et al.*, 2013], match-mismatch dynamics with planktonic food availability [*Frederiksen et al.*, 2006], temperature [*Planque and Frédou*, 1999; *Du Pontavice et al.*, 2022], among other factors [*Myers*, 1998]. However, incorporating environmental drivers of recruitment into stock assessment models can lead to multiple uncertainties, including difficulties identifying recruitment-environment causality, variation in the strength of environmental effects over time, and environmental measurement error where the environmental variable is observed with noise, either from sampling variability or mismatches between what is measured and the environment experienced by the population (e.g. using sea surface temperature as a proxy for temperature at depth). Climate change exacerbates the situation where environmental variables and recruitment-environment relationships are non-stationary, meaning that historically observed data may not be representative of future relationships [*Walters*, 1987].

State space models are an emerging ‘next-generation’ statistical framework within fisheries stock assessment that can account for multiple sources of uncertainty including those associated with environmental effects and underlying environmental processes [*Stock and Miller*, 2021]. The probabilistic structure of state space models allows for a flexible description of model error in the form of random effects - i.e. unspecified stochastic deviations from an otherwise deterministic model structure. This flexibility can be exploited to model uncertainty and variability in environmental relationships and thereby provides a potential path forward to better incorporate environmentally-driven recruitment-environment relationships within fisheries stock assessment models.

Here we conducted a large simulation-estimation study to evaluate the performance of environmentally-driven recruitment relationships in state space assessment models, specifically investigating a recently developed state space assessment framework, the Woods Hole Assessment Model [WHAM *Stock and Miller*, 2021]. We construct 576 operating models (OMs) that differ in key parameters defining the population dynamics and survey quality. We fit five estimating models (EMs) that differ in i) whether an environmental effect is included in the fit; and ii) its functional form. Results from this work will establish the conditions under which we can expect reliable inference from environmentally-driven recruitment relationships in state space assessment models. In doing so, we help advance the effort to better integrate environmental drivers into fisheries management and thereby help prepare management for the increasing impacts of climate change on marine ecosystems.

## Methods

All work described here is based on the Woods Hole Assessment Model (WHAM) [*Stock and Miller*, 2021]. WHAM is an R package freely available as a GitHub repository. All results from this study were produced using WHAM version 1.0.6.9, commit # 77bbd94. WHAM is an increasingly popular state space assessment model now used for management of haddock, butterfish, American plaice, bluefish, Atlantic cod, and black sea bass in waters of the Northeast United States [*NEFSC*, 2022a, b, 2023; *NEFSC*, 2024].

We completed a large simulation-estimation study where OMs differ in the recruitment random effects standard deviation, recruitment random effects autocorrelation, fishing history, environmental autocorrelation, the strength of the environmental relationship, the functional form linking recruitment to the covariate, and the degree of observation error on aggregate survey indices and age composition of both the catch and surveys. See Table S1 for definitions, symbols, and levels used to describe OM factors. OM factor levels were determined, in part, by a review of the range of estimates from recent applications of WHAM for stocks in the Northeast US (Table S2). We fitted five EMs to observations from each of 100 OM model simulations where stochastic process errors were also simulated using a random number generator. EMs made alternative assumptions about the functional form of the environmental driver and/or whether an environmental covariate was used in the fit. Note we did not use the bias-correction feature for process errors or observations described by *Stock and Miller* [2021] for OMs and EMs. Simulations and model fitting were all carried out on the Massachusetts Institute of Technology Green High-Performance Computing Cluster. All code we used to perform the simulation study, summarize results, and generate figures can be accessed publicly at https://github.com/timjmiller/SSRTWG/Ecov_study/recruitment_functions.

### Operating Models

Many of the characteristics of the exploited population in our operating models are identical to those in *Miller et al.* [In review]. Briefly, the population model tracks 10 age classes: ages 1 to 10+ and we assume spawning occurs 1/4 of the way through the year. The maturity at age was a logistic curve with *a*_50_ = 2.89 and slope = 0.88 and assumed known in all EMs. All factors and their levels are given in Table S1. Weight at age *a* was generated with a von Bertalanffy growth function

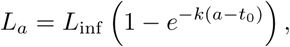

with *t*_0_ = 0, *L*_inf_ = 85, and *k* = 0.3. The length-weight relationship is

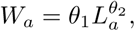

with *θ*_1_ = *e^−^*^12.1^ and *θ*_2_ = 3.2.

Three alternative fishing histories were imposed. In the first scenario, the stock experiences overfishing at twice the fishing mortality rate for maximum sustainable yield (*F* = 2*F*_MSY_) for the first 20 years and *F* = *F*_MSY_ for the last 20 years. In the second scenario, the stock is fished at *F* = *F_MSY_* for the entire time period. In the third scenario, the stock is underfished for the first 20 years at *F* = 0.5*F*_MSY_ then fished at *F* = *F*_MSY_ for the second 20 years. Initial populations were configured at the equilibrium distribution fishing at *F* = 2*F*_MSY_, *F* = *F*_MSY_, or *F* = 0.5*F*_MSY_ depending on the fishing history imposed.

We assumed a single fleet with year-round fishing. Fishing mortality at age was the product of a fully-selected rate and selectivity as a logistic function of age with *a*_50_ = 5 and slope = 1. Two survey time series are simulated in terms of numbers of fish. One survey occurs in the spring and the second in fall. Catchability of both surveys is assumed to be 0.1. Selectivity of both indices was also a logistic function of age with *a*_50_ = 5 and slope = 1, Alternative assumptions about logistic selectivity parameters were also used but results were insensitive to the choice.

Observations of aggregate catch and indices were log-normally distributed and corresponding age composition observations were generated with a logistic-normal distribution with uncorrelated structure on the multivariate-normal scale. Observation error on aggregate indices and the proportion at age (for indices and catch) were tested at a ‘low’ or a ‘high’ level. At the ‘low’ level, the standard deviation of log-aggregate indices was *σ_Iobs_* = 0.1 and and standard deviation parameter for each age of the logistic normal was *σ_paa obs_* was 0.3. At the ‘high’ observation error level, *σ_Iobs_* was 0.4 and *σ_paa obs_* was 1.5. In all scenarios, observation error on aggregate catch *σ_Cobs_* was 0.1, reflecting the convention that catch often approaches a near census and is therefore relatively precise, while age sampling of the catch is more limited and therefore susceptible to measurement error.

All OMs assume a Beverton-Holt (BH) stock-recruitment function with 3 functional forms describing environmental influence (‘none’, ‘controlling’, or ‘limiting’). When there is no environmental driver included, recruitment is given by

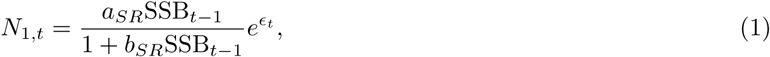

where *a_SR_* and *b_SR_* are the density-independent and -dependent recruitment parameters, respectively, and *ɛ_t_* are first-order autoregressive normal deviations, with parameters described below. We drop the *SR* subscript in the following and allow context to determine whether *a* refers to age or the stock-recruitment parameter. When simulating data from each OM, annual lognormal recruitment deviations are generated with autocorrelation (*ρ_NAA_*) fixed at 0.2 or 0.8. The standard deviation of the log-recruitment deviations varies with values *σ_R_* ∈ {0.1, 0.3, 0.5, 1.0}. We specified unfished recruitment at *R*_0_ = *e*^10^ and *F*_MSY_ = *F*_0%_ = 0.348 which corresponds to a steepness of 0.69, *a* = 0.60, and *b* = 2.4 ∗ 10*^−^*^5^. We note that *F*_MSY_ and *SSB_MSY_* may be time-varying when a time-varying covariate is affecting the stock-recruit relationship. We therefore define *F*_MSY_ and *SSB_MSY_* as the value when the covariate or its effect are 0. Natural mortality is fixed at 0.2. The other two forms that include a driver are given below in the section ‘Environmental link function’.

### Environmental driver

We drive recruitment with a simulated environmental covariate assumed to follow a first-order autoregressive process with zero mean. The conditional form of the process which gives the interannual random effects is given as

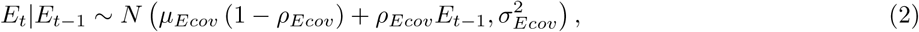

with the marginal (unconditional) form of the process given by

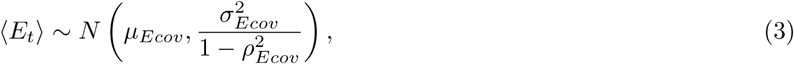

where *E_t_* is the simulated covariate at time *t*, *µ_Ecov_* is the marginal mean, *ρ_Ecov_* is the autocorrelation of *E_t_* and *E_t_*_+1_ which is fixed at 0.2 or 0.8, and *σ_Ecov_* is the conditional random effects standard deviation fixed at 0.1. The annual covariate observation is a normal random variable with mean *E_t_* and we set the observation error standard deviation to 0.1.

### Environmental link function

All OMs assumed a BH functional form. A subset of OMs included the environmental link. The data simulated from all OMs were fit by EMs that specified either a BH or a mean recruitment model. The model for environmentally driven mean recruitment where no stock recruitment function is assumed is given as

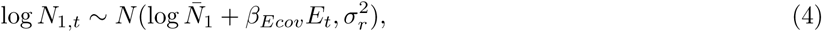

where *N*_1*,t*_ is the number of age 1 recruits in year *t*, log *N̅*_1_ is the mean recruitment on the log scale, *E_t_*is the covariate in year *t* and *β_Ecov_*is the covariate effect size.

OMs with environmentally driven BH functions can take two functional forms, ‘controlling’ or ‘limiting’. These two forms of environmental influence on recruitment were first described in *Fry* [1949] and their parameterization in the BH function were derived in *Iles and Beverton* [1998]. In the controlling functional form, density-independent survival of recruits is modified by an environmental factor, given as

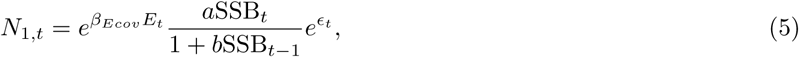

where SSB*_t_* is the spawning stock biomass in year *t*. In the ‘limiting’ functional form, the environment affects the carrying capacity, given as

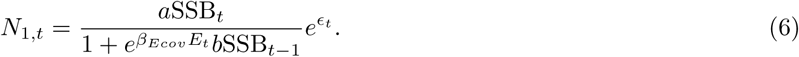

### Estimating Models

We fit five EMs to each of the 100 stochastic simulations across all 576 OM configurations. EMs differed according to the recruitment functional form and whether or not an environmental covariate was used in the fit. All three functional forms were used in the EMs (mean recruitment, controlling, and limiting) with a simulated environmental driver. We also fit mean recruitment and BH recruitment without an environmental driver, totaling five EMs. Both controlling and limiting functional forms reduce to the standard BH when the environmental driver is removed.

### Analysis

We characterized identifiability and performance via a series of statistical diagnostics of the fitted results. We quantified identifiability as the proportion of times the true OM was selected among the set of five EMs using AIC model selection. For performance, we used relative error (RE) to characterize bias, root mean squared error (RMSE) to characterize accuracy, and Mohn’s *ρ* to assess retrospective patterns. We analyzed estimation performance for model parameters listed in Table S3, along with recruitment, SSB, F, and catch (the latter is only assessed in projections). We examined three time periods: 1) all assessment years with observations; 2) the terminal assessment year with observations; 3) ten retrospective forecast years. For the third time period, we removed the last 10 years of data, fit the model, and then projected to the last observed year. When the EM included an environmental covariate, we forecasted future years of the covariate with three approaches: 1) the terminal year covariate held at its last observed value (denoted ‘Constant’); 2) the mean covariate over the 30 observed years (‘Average’); 3) the true simulated covariate (‘True’).

For all diagnostics, we present marginal averages across OM factors and two derived factors (Table S1), along with two-tier regression trees to identify the first- and second-order conditional relationships explaining the most variance across diagnostics. The regression tree approach was adapted from *Hart and Fay* [2020]. Derived factors include i) the realized linear covariate trend generated from stochastic OM model simulations (Δ*E_cov_*); and ii) the realized coefficient of variation of spawning stock biomass (*CV_SSB_*). In all analyses we first dropped simulations that did not pass a series of convergence checks. A model was considered to have converged if the following were true: 1) the optimization routine (stats::nlminb) completed without error; 2) the stats::nlminb convergence flag = 0 indicated successful convergence; 3) the maximum absolute value of the gradient of the log-likelihood was *<* 10*^−^*^6^; 4) TMB::sdreport provided non-NA values for all fixed effects standard errors; 5) TMB::sdreport provided all standard errors *<* 100. We summarized the number and % of crashes across all OM-EM simulations.

## 1 Results

### 1.1 1. Convergence

Of all OM-EM simulations, 26.5% failed one or more convergence checks (Figure 1). This was most pronounced for simulations with constant fishing pressure at *F*_MSY_ when fitting a BH stock recruitment function. For simulations with decreasing and increasing fishing histories (2*F*_MSY_-*F*_MSY_, 0.5*F*_MSY_-*F*_MSY_), higher *σ_r_*led to a lower convergence rate. Across all cases, simulations using mean recruitment maintained high convergence rates with fewer than 5% failing to converge. When considering convergence for EMs with the lowest AIC, the rate of convergence failure dropped to 8.4%, indicating that the fit with the lowest AIC also occasionally failed at least one criterion (Figure 2). These cases were dominated by simulations with the lowest *σ_r_*. The most common reason for non-convergence was the magnitude of the gradient of the cost function at the optimized parameters. Our threshold of *>* 1 × 10*^−^*^6^ was exceeded most often by the stock recruitment parameters of the BH function (consistent with parameter estimation results below).

**Figure 1:**
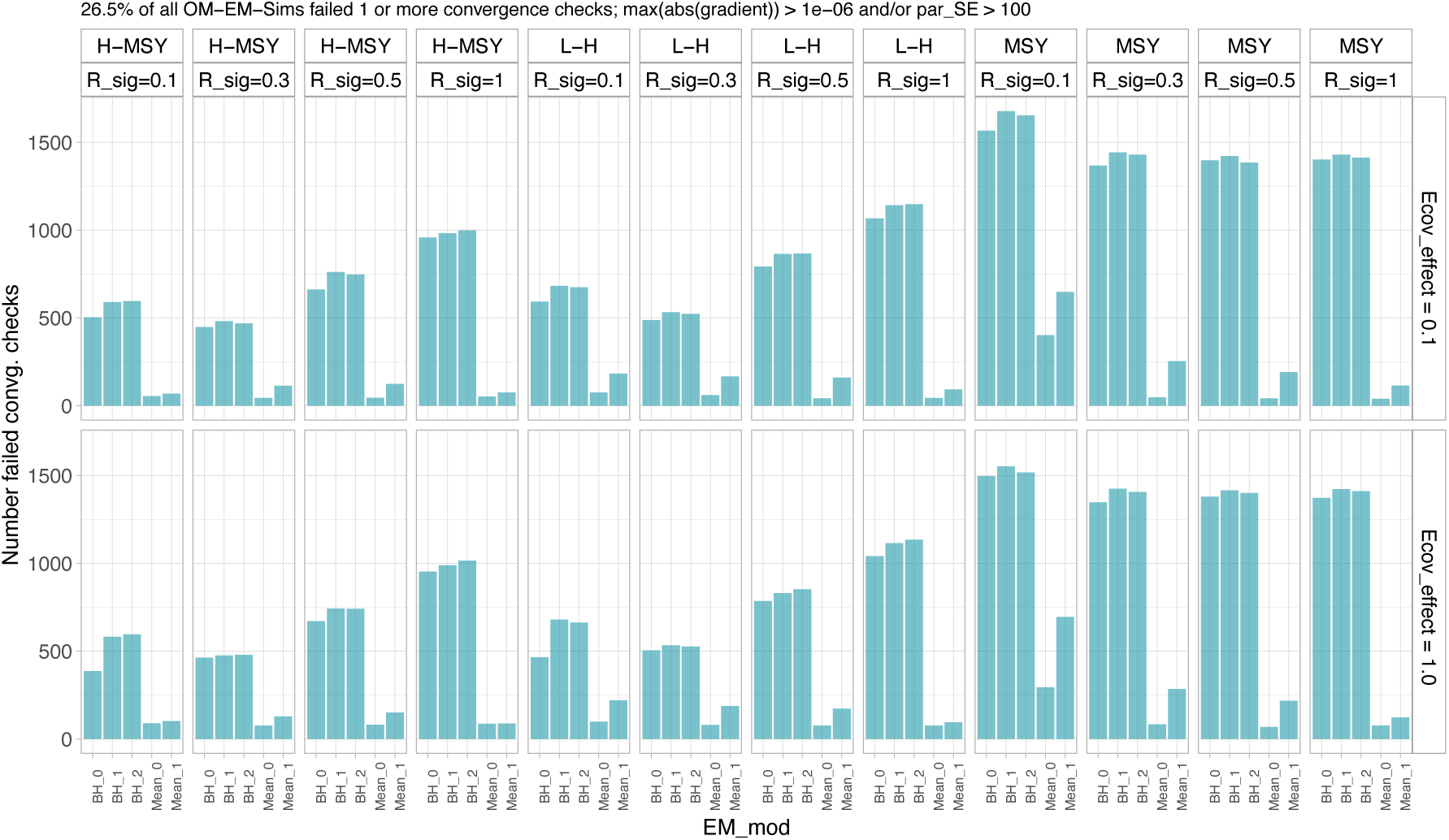
The number of simulations for which the EM failed one or more convergence checks. The x-axis labels give the EM according to whether the Beverton-Holt (BH) or mean recruitment (‘Mean’) functional form is used. If BH=0 or Mean=0 means no environmental link. BH = 1 is the controlling functional form, BH = 2 is the limiting functional form. Figure panels give OM factors that best explained convergence variability.

**Figure 2:**
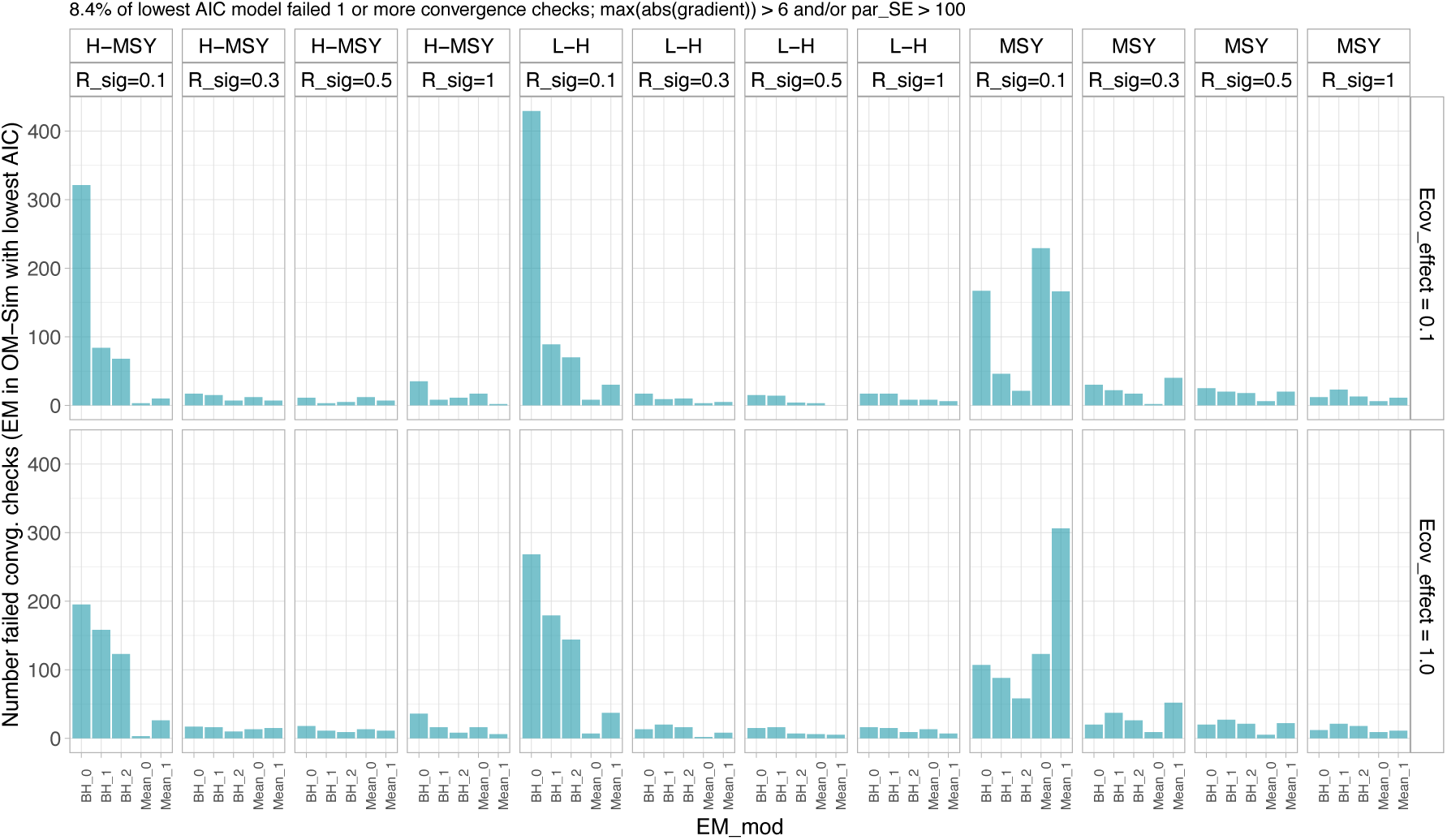
The number of simulations for which the best fitting EM (according to AIC) failed one or more convergence checks. The x-axis labels give the EM according to whether the Beverton-Holt (BH) or mean recruitment (‘Mean’) functional form is used. If BH=0 or Mean=0 means no environmental link. BH = 1 is the controlling functional form, BH = 2 is the limiting functional form. Figure panels give OM factors that best explained convergence variability.

#### Model identification

##### Correct SRR

Model selection results indicated a strong preference for selecting mean recruitment over the BH function. All OMs contained the BH function so this resulted in the correct SR relationship being selected approximately 38% across all OM-EM simulations (Figure 3a). The strongest factor effects were for *σ_r_*, Fhistory, and *CV*_SSB_. Increasing *σ_r_* caused reduced model selection accuracy, while constant fishing pressure (*F* = *F*_MSY_) and consequently low *CV*_SSB_ also caused reduced accuracy. Conditional relationships revealed via regression trees showed that an accuracy of 87% could be achieved with fishing contrast and the lowest *σ_r_* level (*σ_r_*=0.1; Figure S1a).

**Figure 3:**
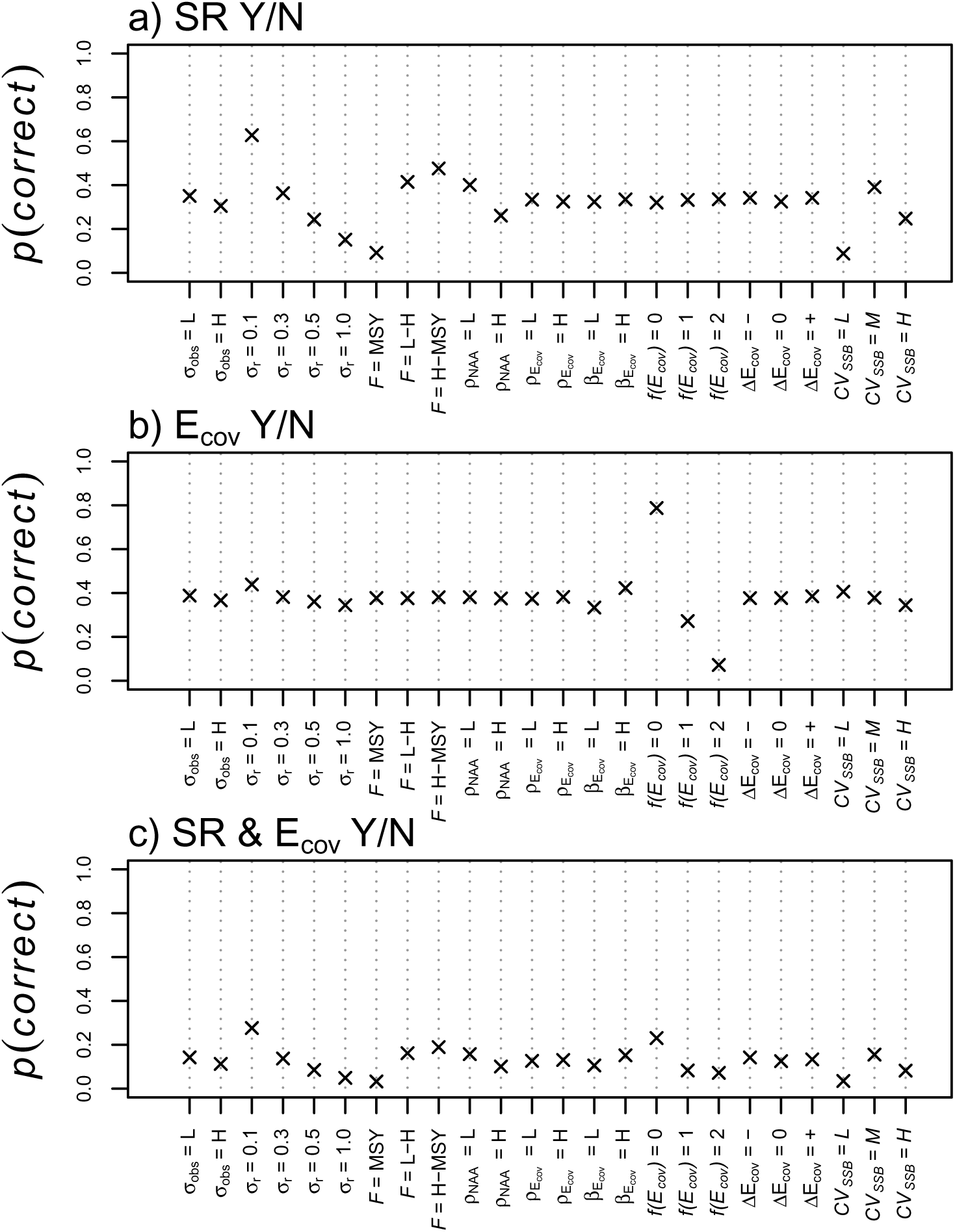
Marginal mean classification accuracy with respect to OM factors. *p(correct)* gives the proportion of OM-EM simulations classified correctly. SR Y/N indicates whether a stock recruitment function was correctly identified (a). E_cov_ Y/N indicates whether the correct functional form of environmental driver was identified (b). SR & E_cov_ Y/N indicates whether an SR relationship and the correct functional form was identified (c). OM factors are defined as in the Table S1

##### Correct environmental mechanism

Mean model selection accuracy for choosing the correct environmental mechanism was similar to that when choosing the BH function overall, achieving just under 40% model selection accuracy across all OM factors (Figure 3b). Individual OM factors played a relatively weak role except for the simulated form of the environmentally-driven stock recruitment function where *f* (*E_t_*) = 0 (no environmental effect) achieves a two-fold higher selection accuracy at approximately 80%. Regression trees again indicate the large difference in accuracy when selecting for an explicit environmental mechanism (Figure S1). The combination of no environmental effect and constant fishing pressure resulted in 80% selection accuracy.

##### Correct SRR and environmental mechanism

Simultaneous identification of an SRR with the correct environmental mechanism was generally low at approximately 18% across OM factors, or slightly less than half of that for identifying the SRR or the correct environmental mechanism alone (Figure 3c). Leading factors driving this result were *σ_r_*, due to variation in SR identifiability, and *f* (*E_t_*) = 0, due to variation in environmental mechanism identifiability. Regression trees again corroborated this finding (Figure S1c).

#### Δ**AIC**

While model identification was generally low, differences in AIC (ΔAIC) between the best and second best model was only ΔAIC ≈ 1.2, on average (Figure 4). Nearly 20% of the time, the second best model was the true OM. No OM factors had an appreciable effect (Figure 4), corroborated by regression trees (Figure S2). The true OM had ΔAIC *<* 2 almost 30% of the time with no covariate, but that dropped to 20% if there was a covariate.

**Figure 4:**
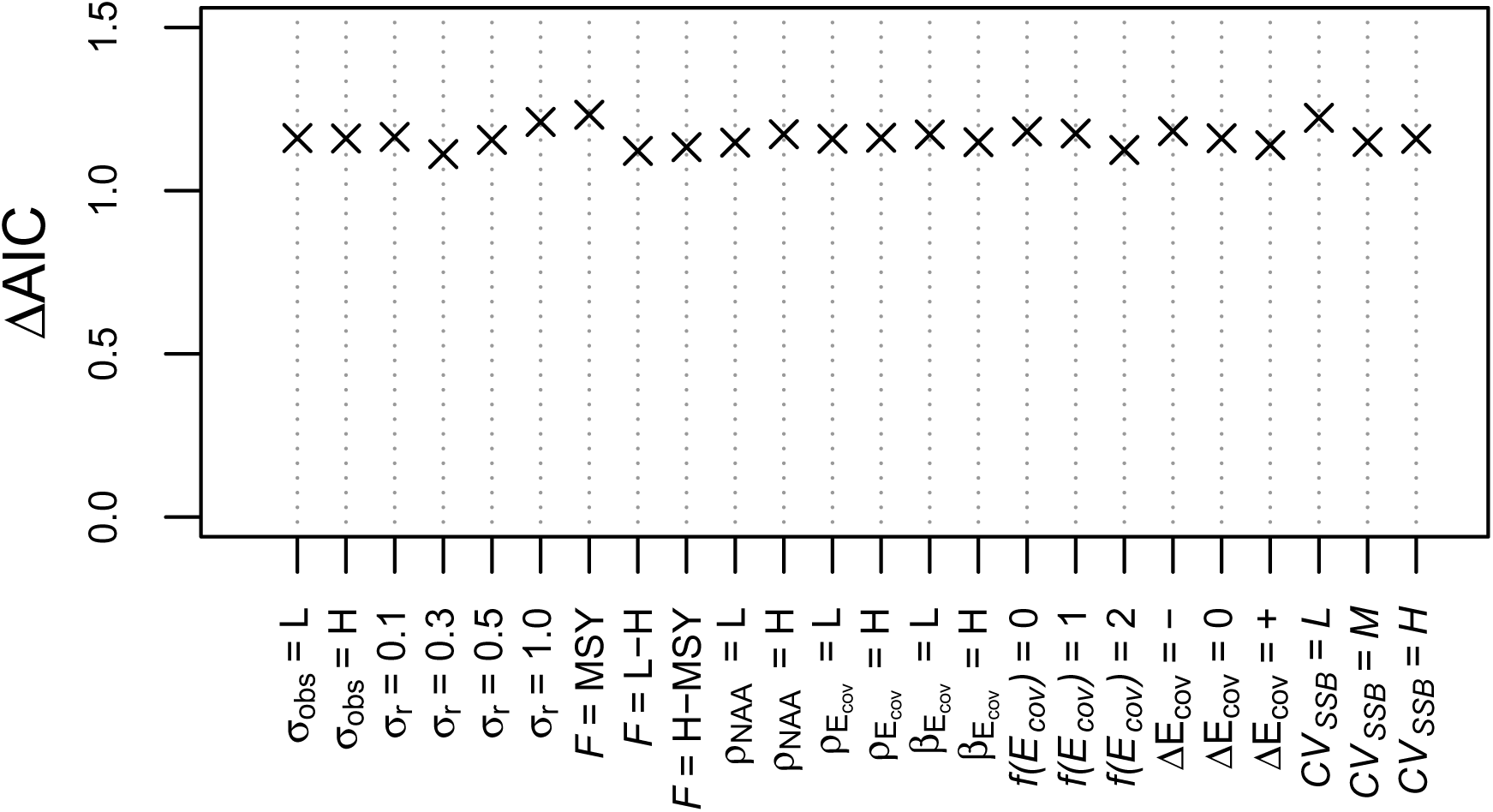
Marginal ΔAIC means averaging across OM factors for the second-best fitting model. OM factors on the x-axis are defined as in Table S1.

#### Parameter estimation performance

Estimation of key model parameters generally appeared unbiased, with a small number of parameters showing considerable spread across OM-EM simulations and a non-zero mean bias (Figure 5). Parameters of note include *σ_r_*, *µ_Ecov_*, *σ_Ecov_*, *a* and *b* (Figure 5f,r,s,u,v). Histograms for these parameters show distributional modes close to zero; however a small number of large biases drive a positive, non-zero bias, on average, particularly for *a*, *b*, and *σ_r_*. Examining relationships between parameter bias and OM factors revealed diverse patterns (Figure 6); most notably, bias in stock recruitment parameters *a* and *b* have a very strong dependence on *σ_r_*, fishing history, and consequently *CV*_SSB_ (Figure 6a,b). Other notable patterns included high parameter bias of *σ_r_* when simulated at the lowest OM level *σ_r_* = 0.1. Bias in the environmental effect *β* also depended strongly on the mechanism of the environmental effect where results showed a high estimation bias for the controlling mechanism (*f* (*E_t_*) = 1) and low estimation bias for the limiting mechanism (*f* (*E_t_*) = 2).

**Figure 5:**
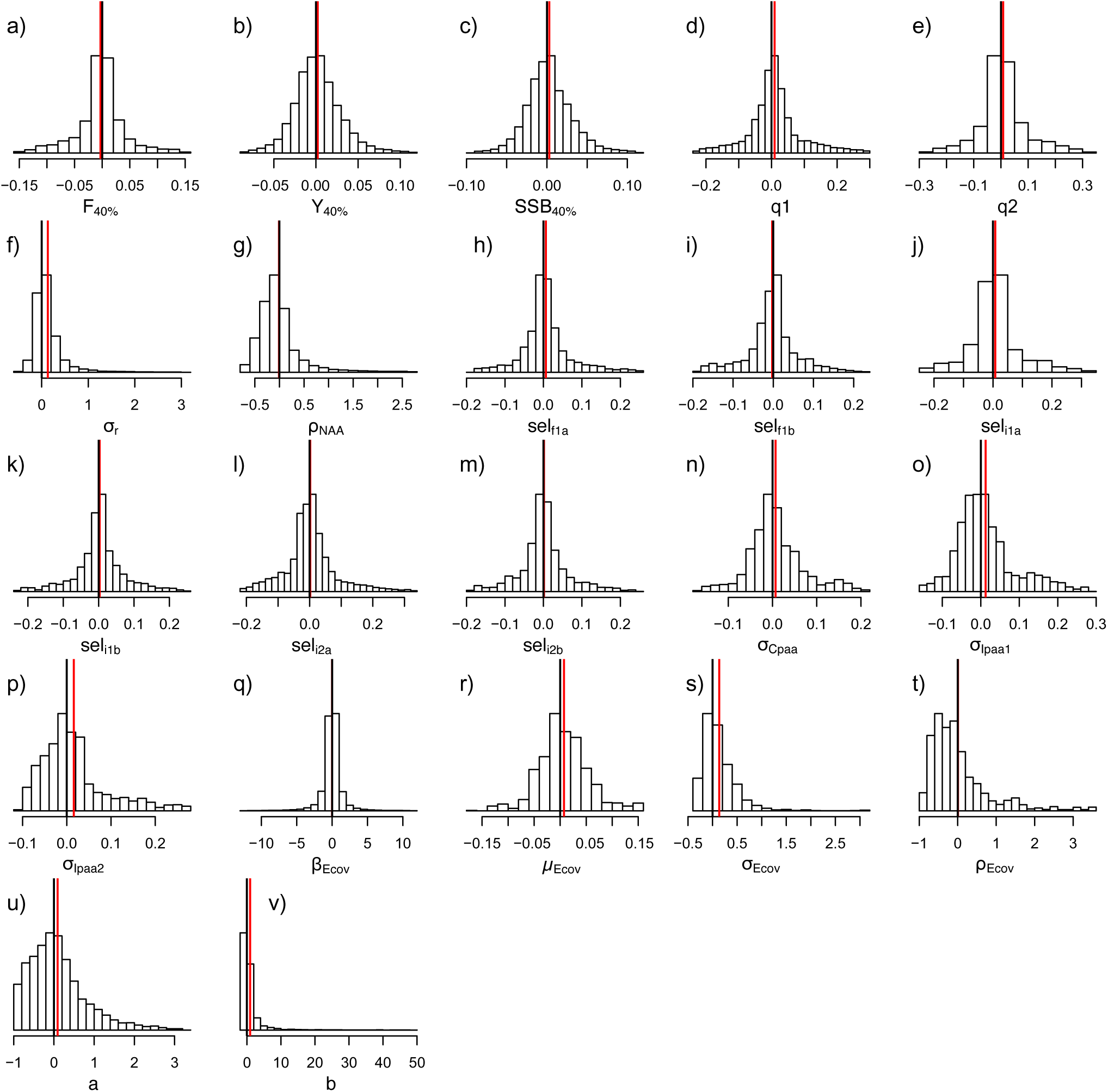
Distribution of relative estimation error for a core set of model parameters. Each parameter is plotted according to 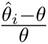, where 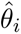 is an individual estimate and *θ* is the true simulated value. Black solid line gives zero line. Red solid line gives mean of the distribution. Model parameters and abbreviations are defined in Table S3.

**Figure 6:**
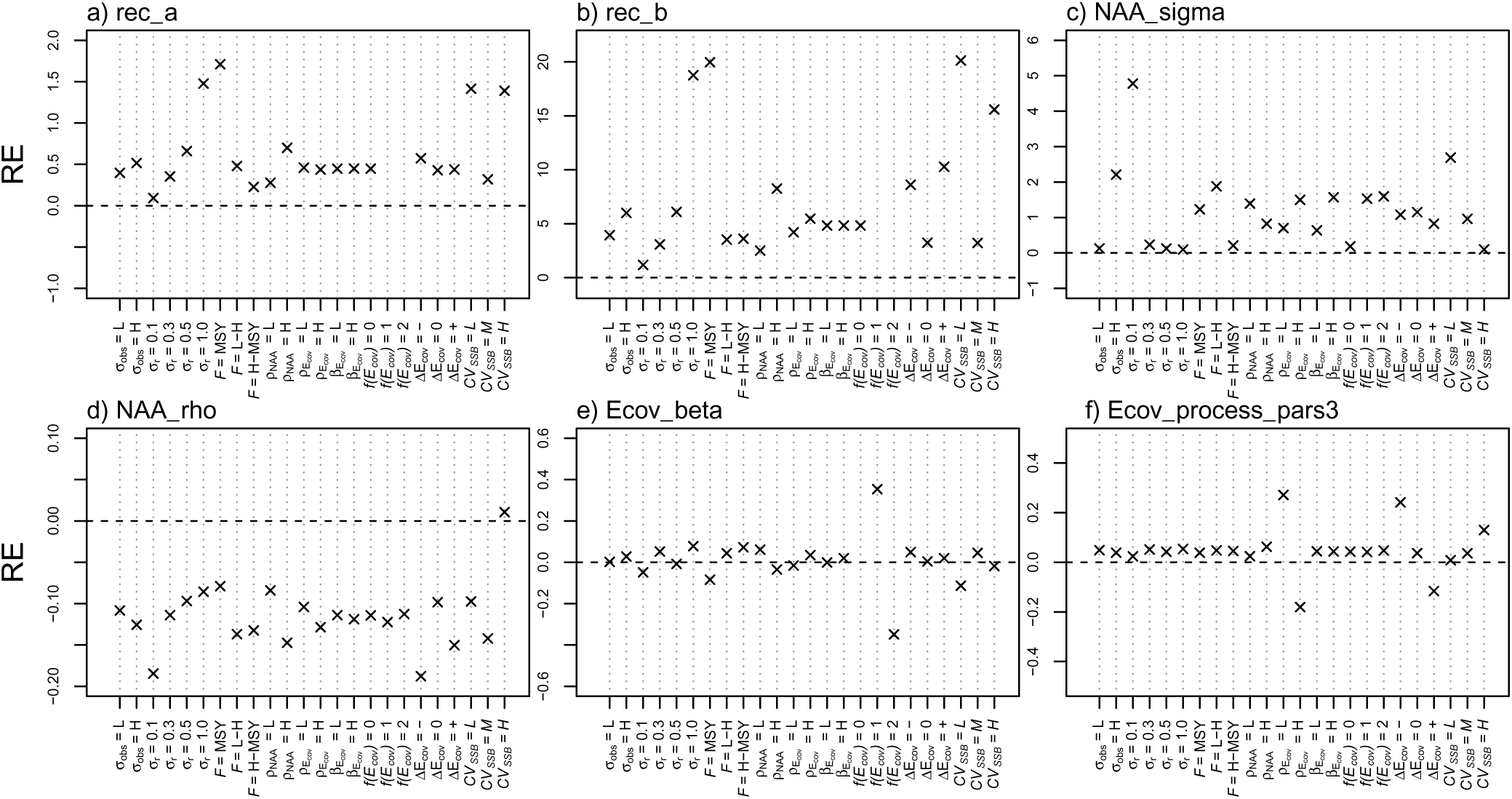
Relative error against OM factors for selected model parameters. OM factors on the x-axis are defined as in Table S1. Model parameters and abbreviations are defined in Table S3.

#### Assessment error in recruitment, SSB, and F

##### All assessment years

Averaging across all assessment years we found mean bias less than 5% in recruitment, SSB, and F, across OM factors (Figure 7a-c). Despite small differences, assessment error was most variable for recruitment, with bias positively related to *σ_r_*and *σ_obs_*. Recruitment and SSB RMSE showed nearly three-fold variation as a function of *σ_r_*(Figure 7g-h). Recruitment and SSB RMSE was also positively associated with the level of realized *CV_SSB_*. Regression trees corroborated the finding that bias remained low across OM factors (Figure S3a-c). Regression trees also showed that *σ_r_*, *σ_obs_*, and *CV*_SSB_ were the most important factors for recruitment and SSB assessment RMSE, respectively (Figure S3d-e).

**Figure 7:**
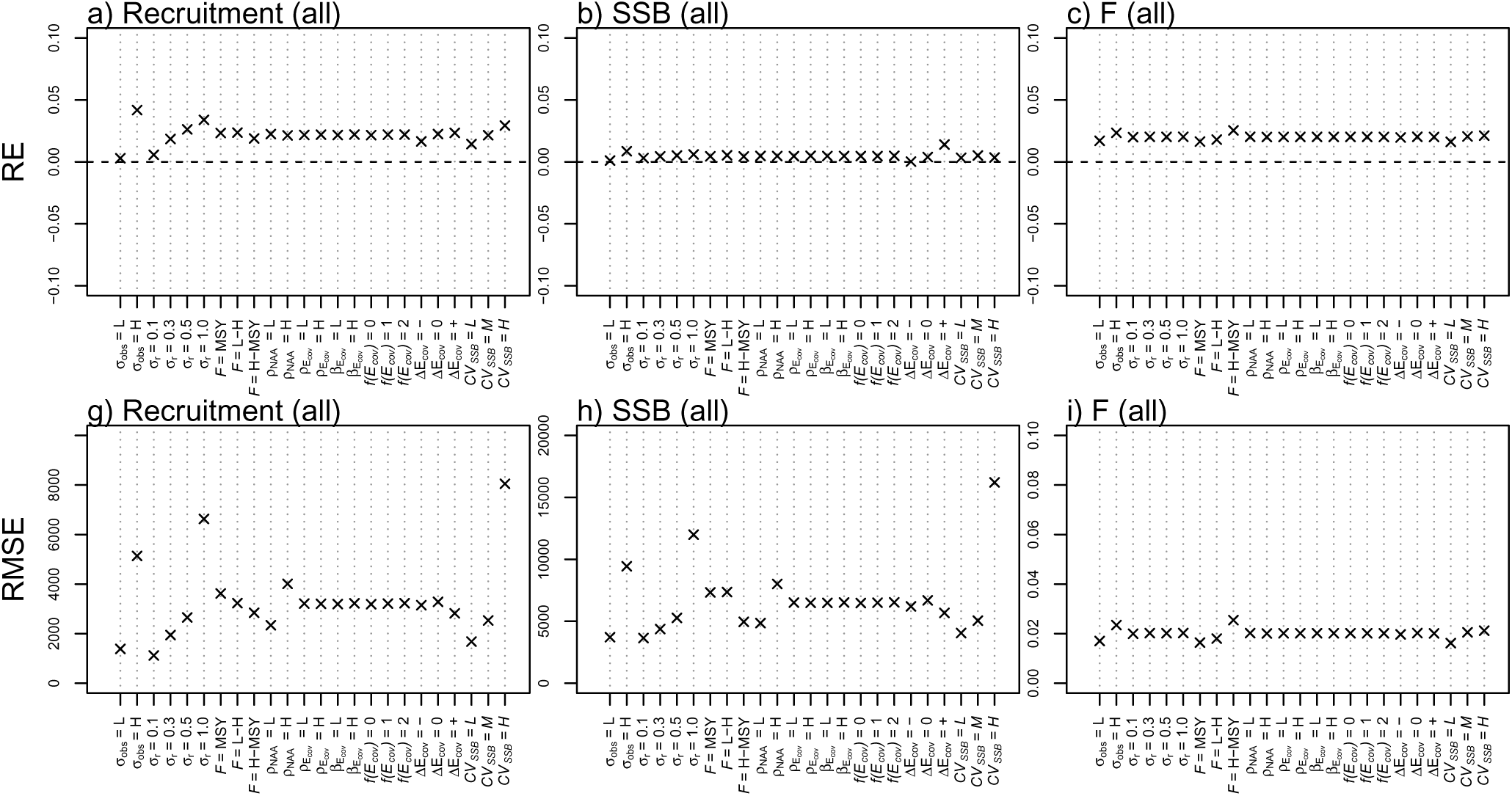
Marginal mean assessment estimation error with respect to OM factors averaged over all assessment years. Columns (left to right) give recruitment, spawning stock biomass (SSB), and fishing mortality (F), respectively. Top row gives relative error (RE). Bottom row gives root mean squared error (RMSE). OM factors on the x-axis are defined as in Table S1.

##### Terminal assessment year

Assessment error in the terminal year was more variable than when averaging over all assessment years, as expected due to statistical averaging. Terminal recruitment and SSB were unbiased across OM factors while F was found to have a slight positive bias of less than 5% (Figure 8a-c). Terminal year recruitment bias was correlated to Δ*E_cov_*and *CV*_SSB_. A positive Δ*E_cov_* led to a negative terminal year recruitment bias while a negative Δ*E_cov_* led to a positive bias. *σ_obs_* and *CV*_SSB_ were positively linked to terminal year recruitment bias. Terminal year SSB and F bias showed less variability with respect to OM factors. Terminal year RMSE for recruitment and SSB was most strongly related to *σ_r_*, *σ_obs_*, and *CV*_SSB_ whereas RMSE for fishing mortality was less variable across OM factors (Figure 8d-f). Regression trees also indicated generally unbiased terminal year assessments but also the importance of Δ*E_cov_* and *σ_obs_*(Figure S4a-c).

**Figure 8:**
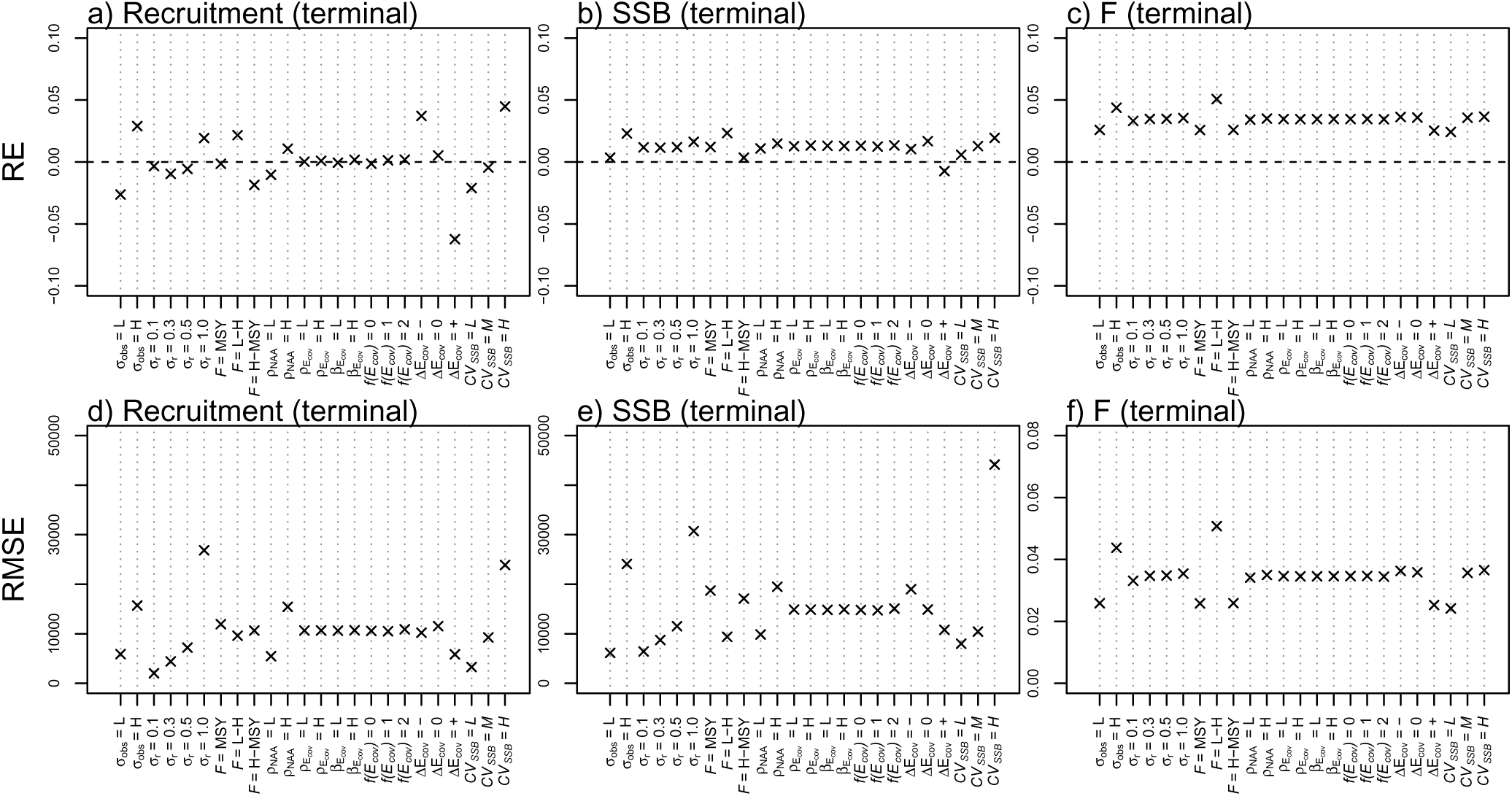
Marginal mean assessment estimation error with respect to OM factors for only the terminal assessment year. Columns (left to right) give recruitment, spawning stock biomass (SSB), and fishing mortality (F), respectively. Top row gives relative error (RE). Bottom row gives root mean squared error (RMSE). OM factors on the x-axis are defined as in Table S1.

#### Retrospective patterns

Averaging across OM factors, mean Mohn’s *ρ* was near zero; however Mohn’s *ρ* for recruitment showed considerable variability with respect to OM factors (Figure 9a). The three main factors driving variability in recruitment Mohn’s *ρ* were *σ_r_*, fishing history, and Δ*E_cov_*. Mohn’s *ρ* was positively associated to *σ_r_*. High fishing history (*F* = *L* − *H*) led to positive values while low fishing history (*F* = 2*F*_MSY_ − *F*_MSY_) led to negative values. The strongest factor was Δ*E_cov_* where Mohn’s *ρ* was negative (−0.04) for negative Δ*E_cov_* and positive (0.04) for positive Δ*E_cov_*. Mohn’s *ρ* was generally constant across OM factors for SSB and F (Figure 9b-c). Regression trees showed that a combination of *σ_r_* = 1 and a positive Δ*E_cov_* could increase Mohn’s *ρ* to a value of 0.12 (Figure S5a), while no pair of OM factors could combine to drive values above/below 6% for either SSB or F (Figure S5b-c). Current guidelines would not consider any of these Mohn’s *ρ* values as significant for management [*Hurtado-Ferro et al.*, 2014].

**Figure 9:**
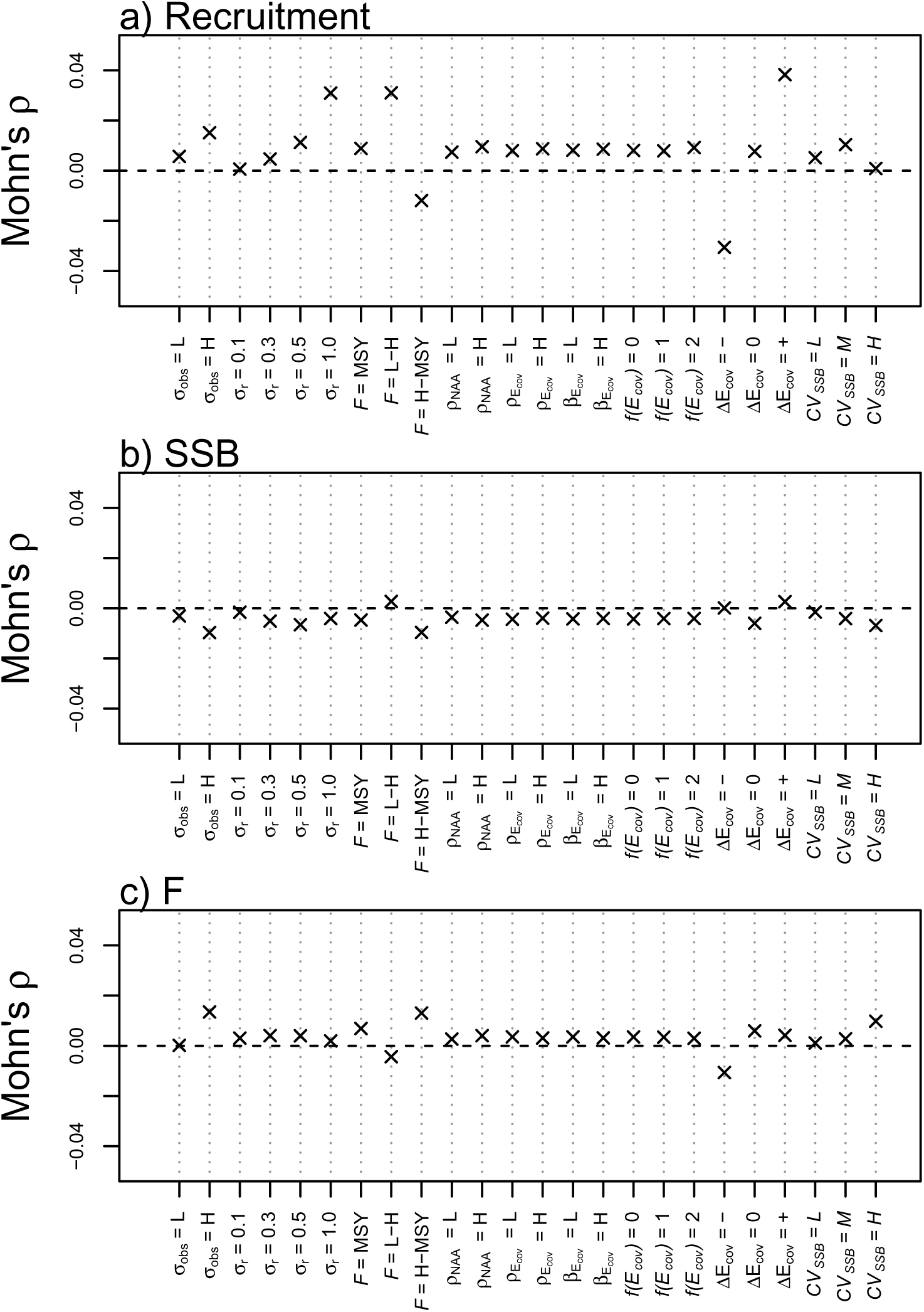
Marginal mean Mohn’s *ρ* with respect to OM factors. Rows (top to bottom) give recruitment (a), spawning stock biomass (SSB) (b), and fishing mortality (F) (c), respectively. OM factors on the x-axis are defined as in Table S1.

#### Projection error in recruitment, SSB, and catch

We found negligible bias in projected quantities and negligible differences between projection scenarios, i.e. where projections i) continued the last observed covariate (‘Constant’), ii) fixed the covariate process at the recent 5 year average (‘Average’), and iii) projections that used the true underlying covariate (‘True’, Figures 10, 11). Examining projection error as a function of time, we find a weak but increasing projection bias with increasing time (Figure 10a,c,e). RMSE also increases with time for SSB and Catch, particularly after 2 or 3 years, but shows more variable patterns for recruitment (Figure 10b,d,f). With respect to OM factors, projection error is nearly identical across ‘Constant’, ‘Average’, and ‘True’ covariate scenarios.

**Figure 10:**
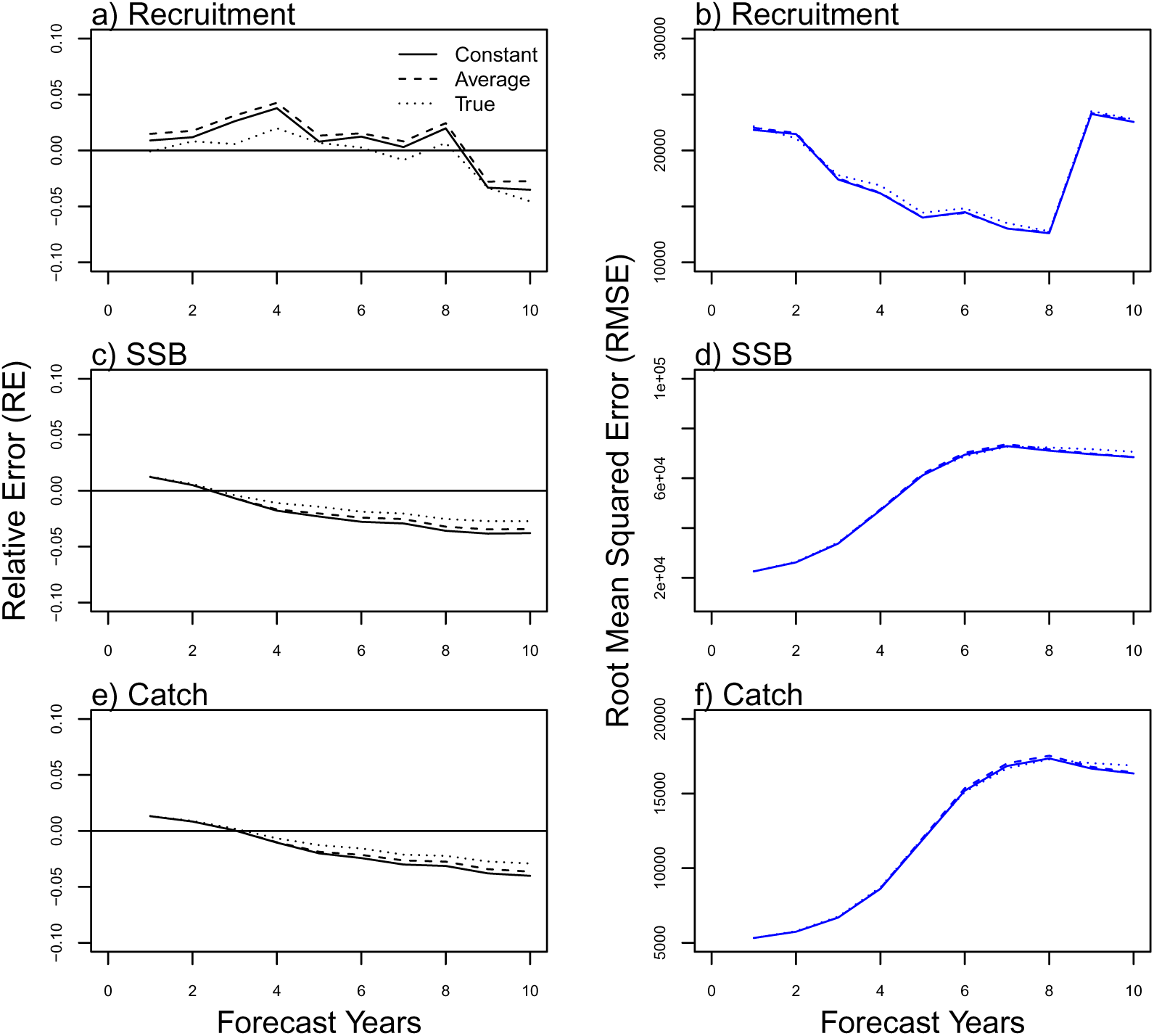
Projection error as a function of time into the future (Forecast Years), averaged over all converged OM-EM simulations. Solid, dashed, and dotted line give cases i) holding the last observed covariate constant “Constant”); ii) averaging all observed covariates (“Average”); iii) using the true underlying covariate (“True”), respectively. Left column gives the relative error (RE). Right column gives the root mean squared error (RMSE). Top, middle, and bottom rows give recruitment, spawning stock biomass, and catch, respectively.

**Figure 11:**
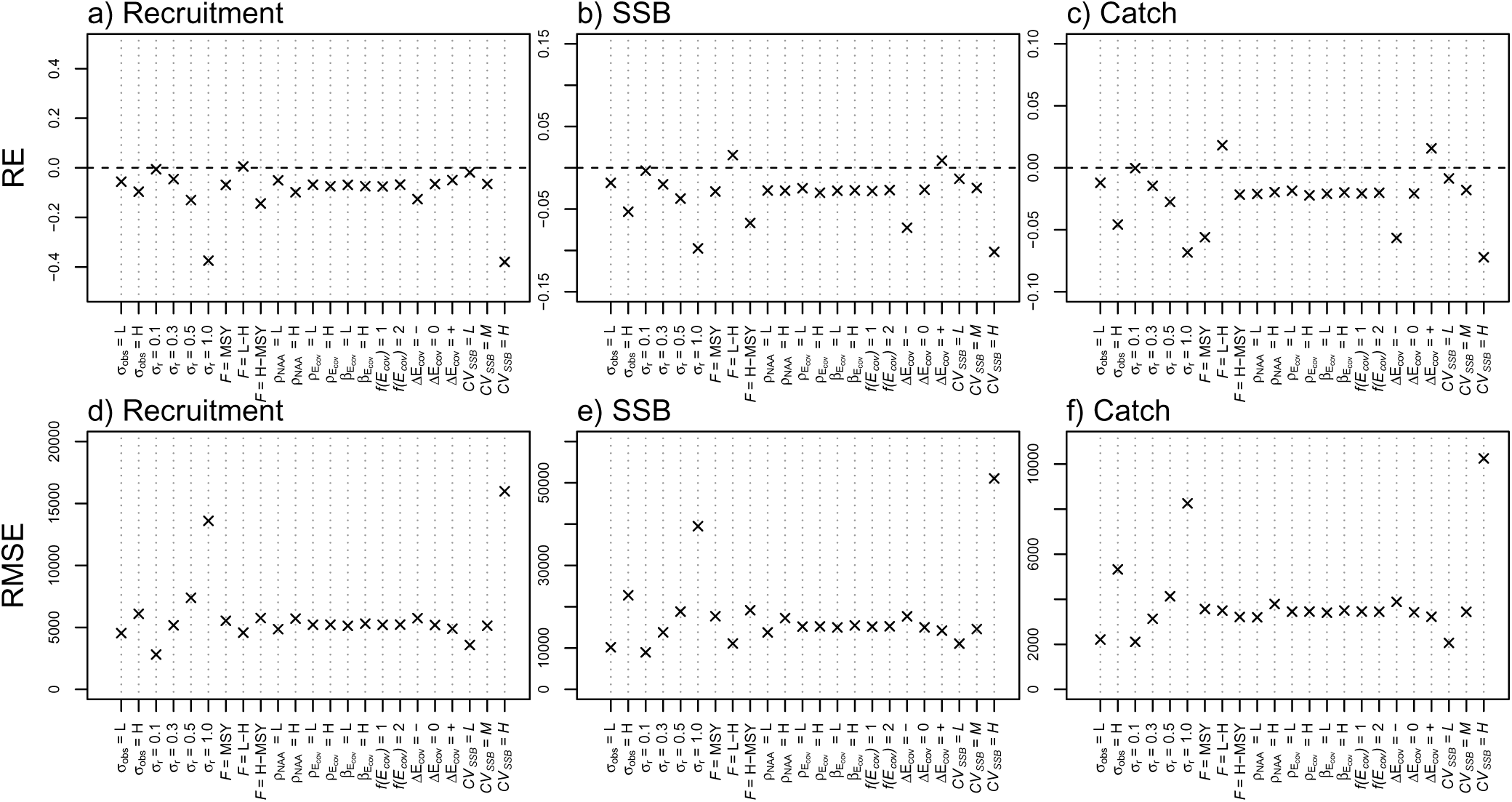
Marginal mean projection error with respect to OM factors. Columns (left to right) give recruitment, spawning stock biomass (SSB), and fishing mortality (F), respectively. Top row gives relative error (RE). Bottom row gives root mean squared error (RMSE). OM factors on the x-axis are defined as in Table S1.

## Discussion

Results presented here help determine the reliability of environmentally-driven recruitment relationships within next-generation state space assessment models, focusing specifically on the increasingly popular WHAM framework. In general we found isolated convergence issues and relatively weak identifiability across simulations. Convergence issues generally appeared for unrealistic values of OM factors (e.g. *σ_r_* = 0.1), while identifiability issues were present across OM factors. Despite poor identifiability using AIC, we generally found low parameter and assessment bias across operating and estimating model combinations, indicating that the state space framework is robust to model mispecification. Robustness to mispecification is a feature of the random effects structure that can adequately account for underlying model errors. Results suggest that Gaussian random effects have an appropriate statistical structure to effectively absorb model errors under the simulated scenarios. The tendency for AIC-based model selection to favor EMs with no SRR (despite a true underlying SRR) also suggests that random effects may allow assessments to avoid unnecessary complexity in estimating an SRR. A similar result was obtained when including an explicit environmental covariate which was correctly identified in only ∼40% of cases but showed no appreciable impact on assessment bias and RMSE, again showing how random effects successfully account for multiple forms of mispecification. Based on these findings, we recommend random effects with no explicit SRR as a simple default model choice in WHAM unless an explicit SRR is favored for other reasons. This reflects current practice and pragmatic guidance [*Skern-Mauritzen et al.*, 2016; *Brooks*, 2024].

In light of other work, we contrast the recruitment random effects approach with methods that apply random effects directly to BH parameters and thereby explicitly resolve the stock-recruitment relationship [*Dorner et al.*, 2008; *Britten et al.*, 2016]. We expect results may be similar when applying random effects to SRR parameters vs. applying them to recruitment deviations, as done here, although we leave those investigations for future work.

We note that SRR parameter random effects have been applied to stock-recruitment time series directly and have not been integrated into a state space assessment framework, although these extensions could be integrated within WHAM in the future.

Perhaps the most surprising aspect of our results was the lack of difference in projection performance when using an explicit environmental covariate. We expected our three scenarios for environmentally-driven recruitment projections (‘Constant’, ‘Average’, and ‘True’) to yield differential predictive accuracy, with simulations using the true underlying covariate to perform best. The True scenario assumes we know what the environment will do into the future, while in reality we would only have access to an uncertain environmental forecast. We expect to see larger differences between these scenarios if we simulated a more extreme environmental effect size as an OM factor. However, observed relationships between fisheries recruitment and individual environmental variables are generally noisy or ephemeral [*Myers*, 1998] and levels of recruitment variability chosen here were in line with other stock assessment models (Table S2). We therefore opted for realistic OM factors over more extreme settings and therefore likely swamped the environmental signal with high levels of recruitment variability and observation error.

A lack of improvement in assessment outcome using more accurate environmental forecasts suggests a poor potential for more accurate environmental forecasting as a means to improve short-term fisheries forecasts [*Haltuch and Punt*, 2011]. This has direct implications for ongoing efforts to improve ecosystem forecasting capacity on annual timescales (but see Haltuch et al. 2019 who suggest that environmental drivers may be successful for species with short pre-recruit intervals or clear bottlenecks in their life history). We did, however, observe an effect of environmental non-stationarity on terminal year assessment bias and Mohn’s *ρ* where a strongly trending covariate results in modest but systematic over- or under-estimated recruitment. This is largely driven by the low information in the data available to estimate terminal year recruitment, causing estimates to shrink to default predictions which includes the covariate effect [*Miller et al.*, In review]. We conclude from these results that environmental predictability in state space stock assessment models may show differences in performance across timescales: On short timescales, recruitment variability and observation error swamp the typical environmental signal resulting in no net assessment improvement. On long timescales, however, non-stationary environmental drivers may cause moderate bias across years and decades, although we note that biases for recruitment may disappear as assessments are updated and more cohort data are available as they age through the fishery. while the magnitude of Mohn’s *ρ* associated with the covariate trend was not large, these preliminary results call for further exploration of non-stationary environmental dynamics, including extensions to WHAM that account for non-stationary environmental time series and environment-recruitment relationships. This extension would allow us to more explicitly test the role of non-stationary covariates in state space assessment performance. WHAM currently does not support trends in the AR(1) dynamics on the covariate, whereas we analyzed the realized Δ*E_cov_* of individual AR(1) simulations over the assessment period.

Temporally varying biological reference points are an important aspect of environmentally driven stock assessments that are not addressed here. We re-iterate that reference points (and therefore stock status) depend on the underlying stock recruitment relationship and current state of the covariate [*Brooks*, 2024]. This generates conceptual hurdles in applying equilibrium concepts underlying maximum sustainable yield to reference points that vary annually within an environmentally driven stock assessment. Related issues arise when a stock can transition from overfished to not-overfished when the environmental driver changes with no associated change in stock biomass [*Szuwalski et al.*, 2023]. Timescales of variation are also important, where managers may desire reference point stability on short timescales but need to accommodate long-term environmental change in decadal projections. These considerations are non-trivial and challenge the fundamental interpretation of biological reference points, which we leave for future work.

While state space models provide a useful tool in characterizing multiple sources of assessment uncertainty, we expect many fundamental aspects of the ‘recruitment problem’ to persist even with widespread adoption of the state space approach. For example, relationships between population and environmental processes remain inherently complex, difficult to identify, and challenging to observe. Furthermore, much of the utility of random effects is not focused on improved predictive accuracy but instead focused on flexibility to account for and propagate multiple forms of model error. In this way, state space models can reduce subjective data weighting decisions [*Punt*, 2017], and also allow managers to more effectively characterize and balance different sources of uncertainty in a fishery and how those processes may change in time.

In summary, the inclusion of random effects within state space fisheries assessment models is now paving the way for a next-generation of models with improved uncertainty quantification and flexible capacity for the integration of environmental drivers. This study focused on the inclusion of environmentally-driven recruitment processes within the WHAM state space assessment framework and showed a robustness of the approach and the utility in using random effects to model sources of variability, despite difficulties in identifying the correct underlying model structure. As impacts from climate change on marine ecosystems accelerate, these results provide a stepping stone to further developments in the ongoing effort to manage fisheries in a changing environment.

## Supplementary Materials

### Supplementary Tables

**Table S1:**
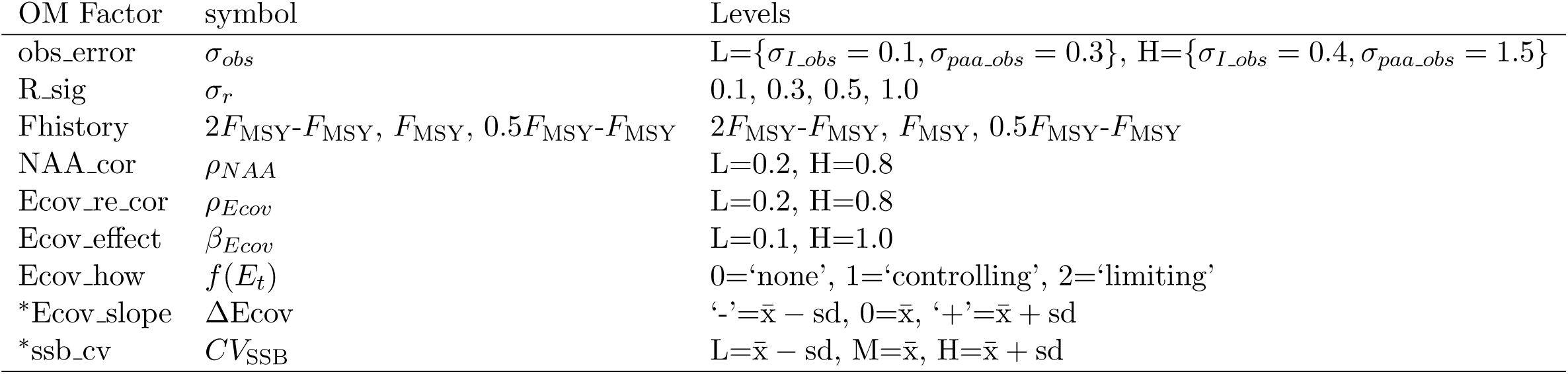
Definitions of operating model factors. Note that Ecov slope and ssb cv are derived quantities that are calculated post-hoc based on properties of the stochastic simulated data. Notation x̄ − sd and x̄ + sd represents intervals defined by one standard deviation from the sample mean.

**Table S2:** Summary of estimates pertaining to random effects estimated for recruitment (*r*) and numbers at age (*NAA*) in recent WHAM assessments. SRR gives the recruitment model used. NAA RE indicates whether random effects were applied to recruitment only (rec) or all numbers at age (rec + 1). RE Cor gives the correlation structure on the random effects. *σ_r_* is defined as in the text. *σ_NAA_* gives the standard deviation of numbers at age beyond the first year class. *ρ_age_* gives the autocorrelation of random effects across ages. *ρ_year_* gives the autocorrelation of random effects across years.

| Stock | SRR | NAA_RE | RE_Cor | $\sigma_r$ | $\sigma_{NAA}$ | $\rho_{age}$ | $\rho_{year}$ |
| --- | --- | --- | --- | --- | --- | --- | --- |
| GB.Haddock | mean | rec+1 | 2dar1 | 1.64 | 0.38 | 0.15 | 0.28 |
| Redfish | BH | rec | iid | 1.40 | NA | 0.00 | 0.00 |
| GB.Cod | mean | rec+1 | iid | 1.25 | 0.52 | 0.00 | 0.00 |
| GM.Haddock | mean | rec+1 | 2dar1 | 1.06 | 0.35 | 0.32 | 0.52 |
| Mackerel | random.walk | rec+1 | 2dar1 | 0.92 | 0.49 | 0.50 | -0.11 |
| N.Black.Sea.Bass | mean | rec+1 | 2dar1 | 0.74 | 0.81 | 0.08 | 0.26 |
| GM.Cod | mean | rec+1 | 2dar1 | 0.68 | 0.25 | 0.34 | 0.70 |
| SNE.Cod | BH | rec | iid | 0.60 | NA | 0.00 | 0.00 |
| Plaice | mean | rec+1 | iid | 0.51 | 0.29 | 0.00 | 0.00 |
| S.Black.Sea.Bass | mean | rec+1 | 2dar1 | 0.51 | 0.60 | -0.13 | 0.33 |
| GB.Winter.Flounder | mean | rec+1 | 2dar1 | 0.34 | 0.19 | 0.63 | 0.69 |
| Bluefish | mean | rec+1 | 2dar1 | 0.33 | 0.16 | -0.20 | 0.77 |
| Butterfish | mean | rec+1 | ar1 | 0.30 | 0.25 | 0.00 | 0.18 |

**Table S3:**
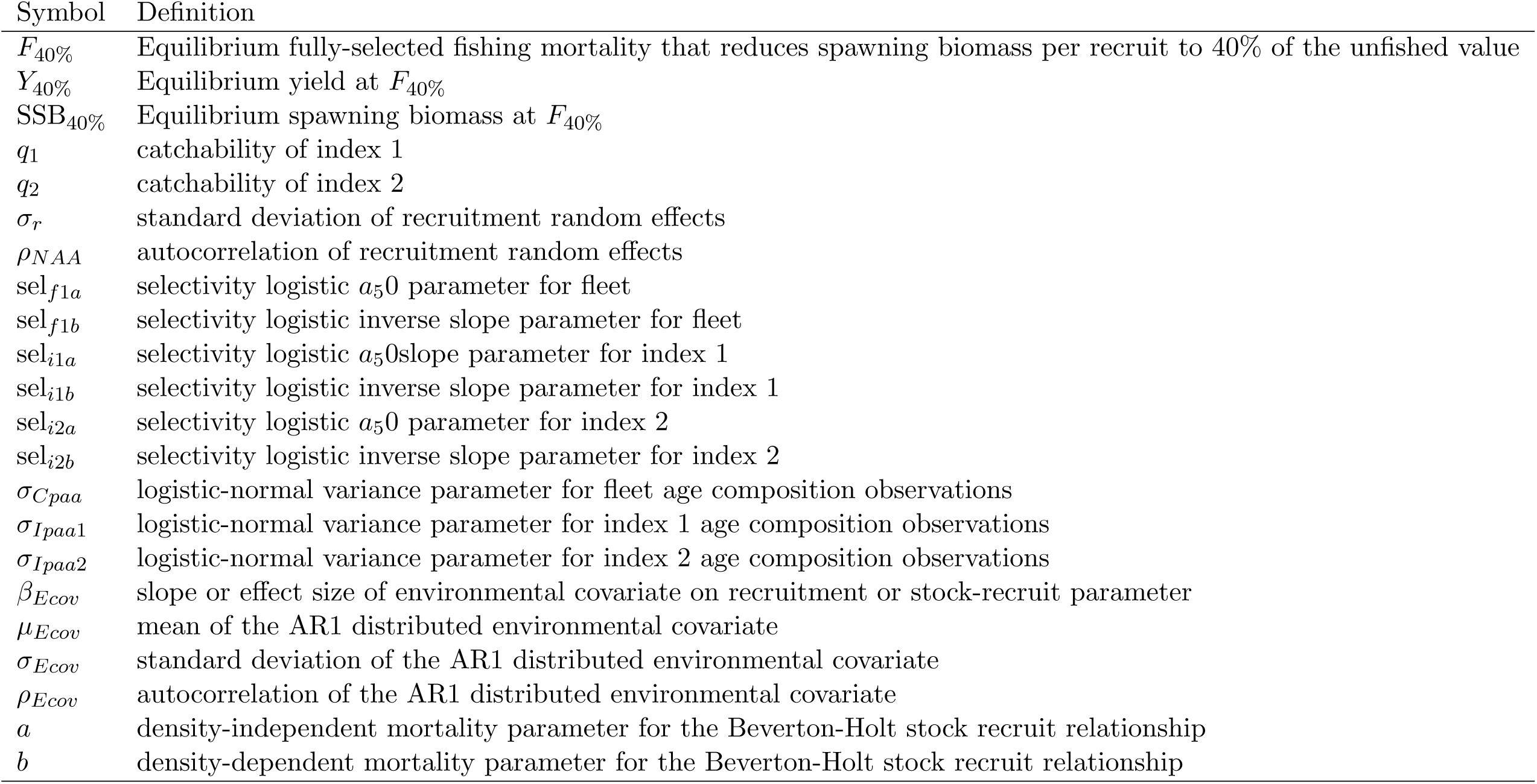
Definitions of model parameters described in the text.

### Supplementary Figures

**Figure S1:**
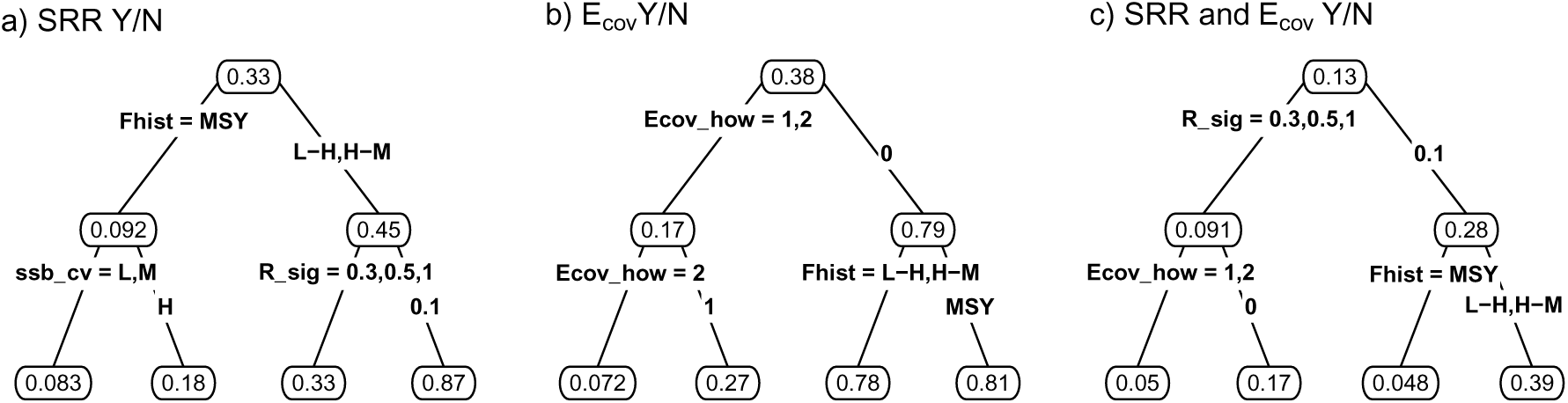
Regression trees of model selection accuracy against OM factors. Nodes are labeled with proportional classification accuracy for that subset. Top node is the overall mean. SRR Y/N indicates whether a stock recruitment function was correctly identified (a). E_cov_ Y/N indicates whether the correct functional form of the SR was identified (b). SRR & E_cov_ Y/N indicates whether an SR relationship and the correct functional form was identified (c). OM factors are defined as in Table S1. Edges are labeled with the factor levels defining the subset.

**Figure S2:**
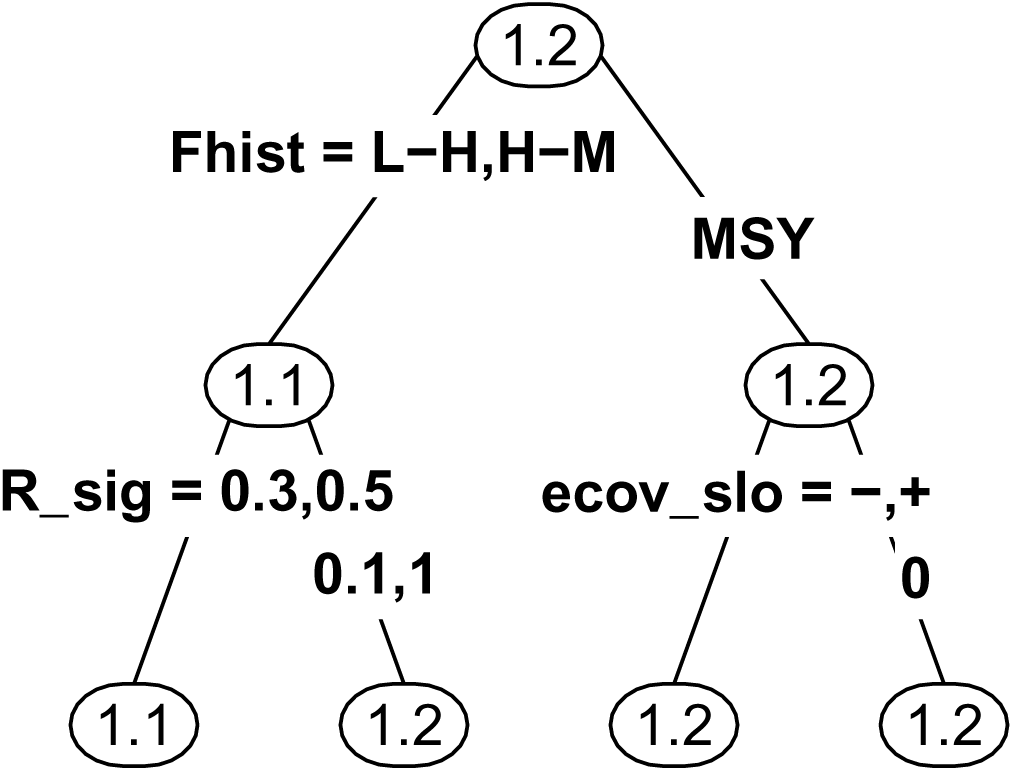
Regression tree of ΔAIC for the second-best fitting model against OM factors. Nodes are labeled with mean ΔAIC for that subset. Top node is the overall mean. OM factors on the x-axis are defined as in Table S1. Edges are labeled with the factor levels defining the subset.

**Figure S3:**
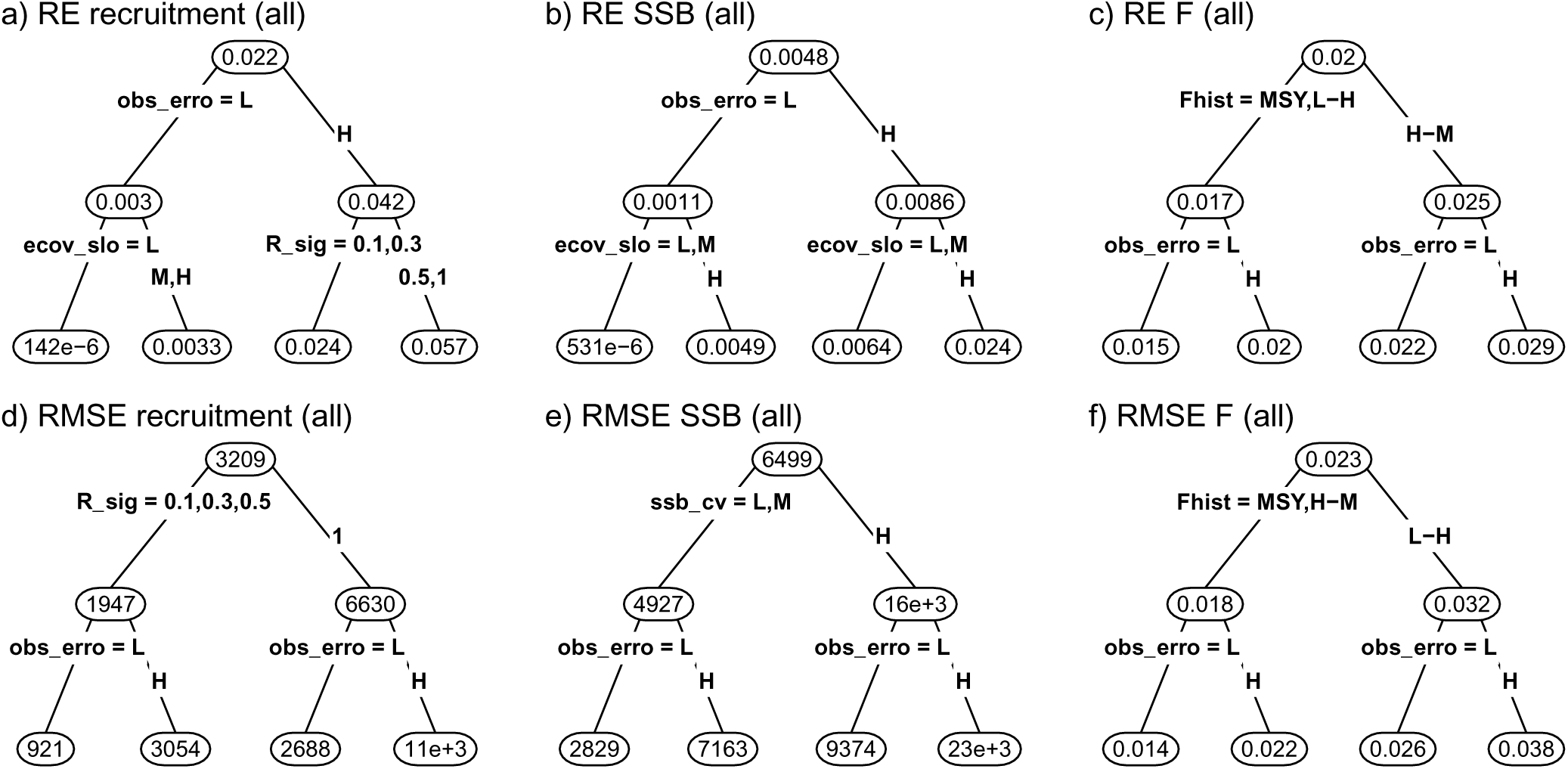
Regression trees of assessment estimation error with respect to OM factors averaged over all assessment years. Columns (left to right) give recruitment, spawning stock biomass (SSB), and fishing mortality (F), respectively. Top row gives relative error (RE). Bottom row gives root mean squared error (RMSE). Nodes are labeled with the mean RE or RMSE for that subset. Top node is the overall mean. Edges are labeled with the factor levels defining the subset as defined in Table S1.

**Figure S4:**
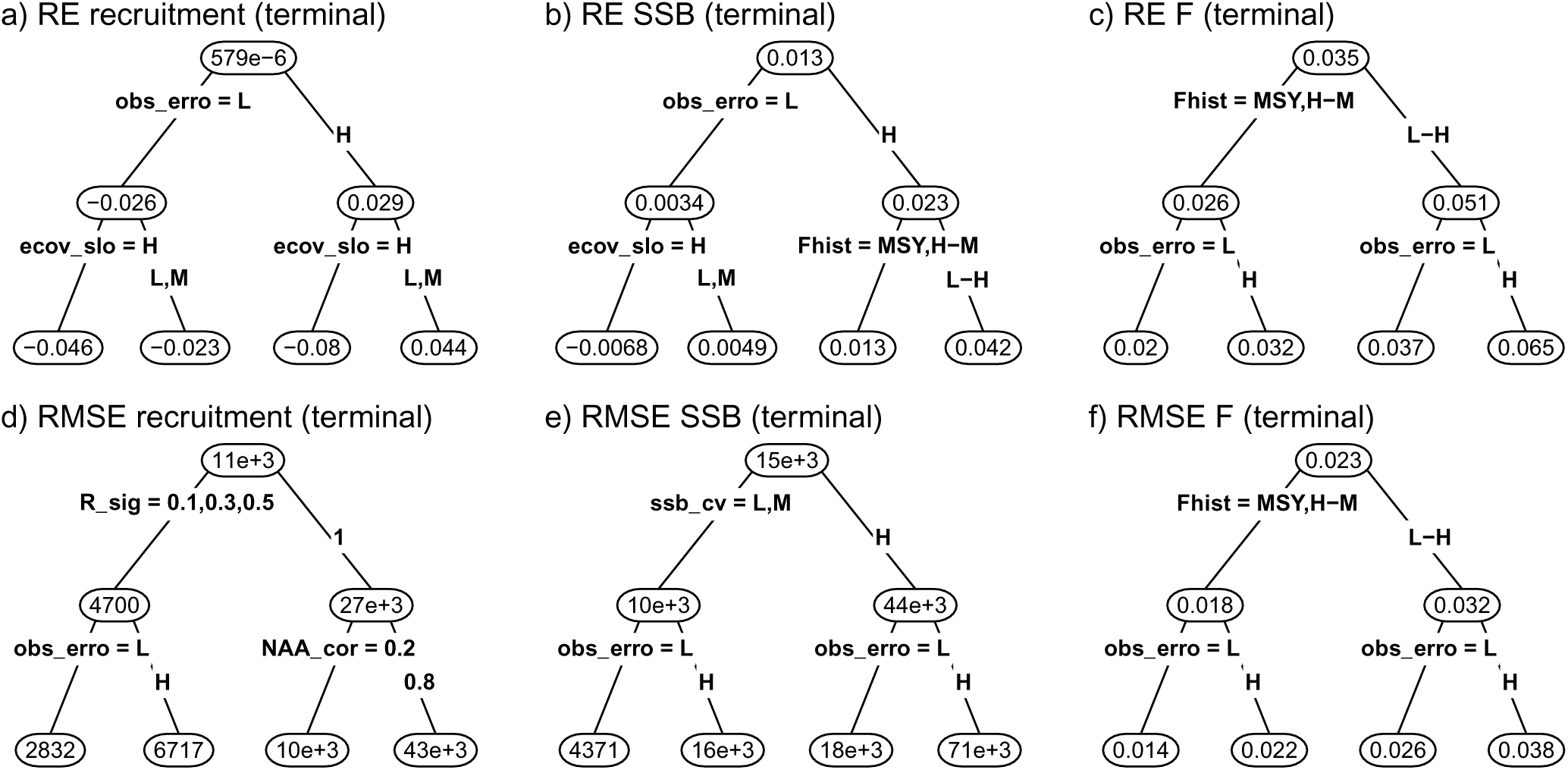
Regression trees of assessment estimation error with respect to OM factors for only the assessment year. Columns (left to right) give recruitment, spawning stock biomass (SSB), and fishing mortality (F), respectively. Top row gives relative error (RE). Bottom row gives root mean squared error (RMSE). Nodes are labeled with the mean RE or RMSE for that subset. Top node is the overall mean. Edges are labeled with the factor levels defining the subset as defined in Table S1.

**Figure S5:**
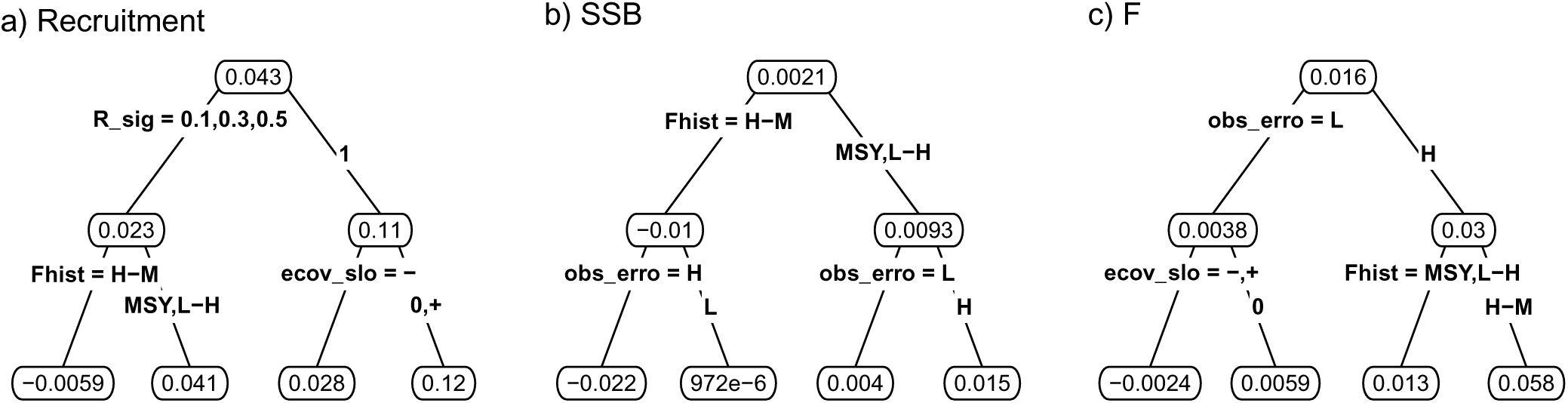
Regression trees of Mohn’s *ρ* with respect to OM factors. Left to right give recruitment (a), spawning stock biomass (SSB) (b), and fishing mortality (F) (c), respectively. OM factors on the x-axis are defined as in Table S1.

